# Identification of a RIPK2-Regulated Gene Signature as a Candidate Biomarker for RIPK2 Activity and Prognosis in Prostate Cancer

**DOI:** 10.1101/2025.04.30.651490

**Authors:** Ahmed M. Elgehama, Qian Yang, Zaoke He, Lauryn Ruegg, Sungyong You, Wei Yang

## Abstract

Receptor-interacting protein kinase 2 (RIPK2) has emerged as a promising drug target in various cancers, including prostate cancer (PC). However, the absence of reliable biomarkers to assess RIPK2 activity limits both patient selection for anti-RIPK2 therapies and treatment monitoring. To address this gap, we performed RNA-Seq analysis on PC cell lines (22Rv1, DU145, and PC3) with CRISPR/Cas9-mediated *RIPK2* knockout (*RIPK2*-KO) using two independent guide RNAs. This analysis identified 13 candidate RIPK2-regulated genes, of which eight were validated by reverse transcription quantitative PCR (RT-qPCR). Furthermore, treatment with two distinct RIPK2 inhibitors significantly reduced RIPK2 signature scores in five independent PC cell lines in a dose- and/or time-dependent manner. Clinical association analyses revealed that high RIPK2 signature scores correlate with metastasis and worse biochemical recurrence-free, progression-free, disease-free, and overall survival, outperforming RIPK2 mRNA levels as a prognostic biomarker. This study establishes, for the first time, a RIPK2-regulated gene signature as a potential biomarker for RIPK2 activity and PC prognosis, warranting further validation in clinical specimens to provide a much-needed tool for patient stratification and response monitoring in RIPK2-targeted therapies.

## INTRODUCTION

Receptor-interacting protein kinase 2 (RIPK2) is a Ser/Thr/Tyr kinase best known for its role in inflammation and innate immunity, primarily through the NOD/RIPK2/NF-κB signaling pathway [1]. Recently, RIPK2 has also been implicated in cancer, where its overexpression across multiple cancer types correlates with disease progression and poor prognosis [2, 3]. In response, many RIPK2 inhibitors have been developed, with several advancing into early-stage clinical trials [1, 4].

Prostate cancer (PC) remains a major health concern, particularly in the United States and other developed countries [5, 6]. With an aging population, global PC mortality is projected to rise to nearly 700 000 deaths annually by 2040, underscoring an urgent need for novel therapeutic strategies [7]. Notably, most of these deaths result from metastasis, which is associated with poor prognosis and significant deterioration in quality of life.

Our recent work identified RIPK2 as a key driver of PC progression and metastasis through a non-canonical RIPK2/MKK7/c-Myc phosphorylation pathway [2]. Targeting RIPK2 effectively suppressed PC metastasis *in vitro* and *in vivo*, positioning it as a promising therapeutic target for advanced PC [2]. However, clinical translation is hindered by the absence of reliable biomarkers to assess RIPK2 activity, which complicates patient selection and treatment monitoring for anti-RIPK2 therapies.

While RIPK2 protein expression has been considered a potential biomarker, it does not accurately reflect RIPK2 activity. This is because RIPK2 activity is regulated by post-translational modifications, protein-protein interactions, and subcellular localization, rather than protein abundance alone. Therefore, relying solely on RIPK2 protein levels risks misclassifying RIPK2-active tumors. In contrast, a RIPK2-regulated gene signature would capture downstream transcriptional outputs of active RIPK2 signaling, providing a more functional and dynamic readout. Although RIPK2 is a potent activator of c-Myc in PC cells [2], MYC signatures are not specific enough for this purpose, as c-Myc activity is modulated by diverse signaling pathways.

Here, we address this critical gap by identifying a RIPK2-regulated gene signature through RNA-Seq analysis, followed by validation using reverse transcription quantitative PCR (RT-qPCR). We further evaluate its association with PC prognosis, laying the groundwork for future clinical application.

## MATERIALS AND METHODS

### Cell lines, culture, and treatment

The PC cell lines 22Rv1 (#CRL-2505), DU145 (#HTB-81), PC3 (#CRL-1435), C4-2B (#CRL-3315), and MDA-PCa-2b (#CRL-2422) were obtained from the American Type Culture Collection (ATCC, Manassas, VA, USA). Control and *RIPK2*-KO 22Rv1, DU145, and PC3 cells were generated using CRISPR/Cas9 technology [2]. PC3, 22Rv1, and C4-2B cells were cultured in RPMI-1640 medium (ATCC, #30-2001) supplemented with 10% fetal bovine serum (FBS; ATCC, #30-2021) and Gentamicin (25 mg/mL) and Amphotericin B (125 µg/mL) (Sigma, St. Louis, MO, USA, #SBR00045-2ML). DU145 cells were cultured in DMEM medium (ATCC, #30-2002) supplemented with 10% FBS and 1% penicillin/streptomycin. MDA-PCa-2b cells were cultured in BRFF-HPC1 medium (Fisher Scientific, Pittsburgh, PA, USA, #NC9970798) supplemented with 10% fetal bovine serum and 1% penicillin/streptomycin. For drug treatment experiments, cells were treated with ponatinib (MedChemExpress, Monmouth Junction, NJ, USA, #HY-12047), RIPK2-IN-4 (MedChemExpress, #HY-107978), or AZD1480 (VWR International, Radnor, PA, USA, #ALA126326) at the indicated doses and time points.

### RNA sequencing and data analysis

Total RNA was extracted from cells using the RNeasy Mini Kit (Qiagen, Valencia, CA, USA, #74104). RNA sequencing was performed by BGI Group (Shenzhen, China) using the DNBseq platform with a DNBSEQ Eukaryotic Strand-specific mRNA library and PE150 read length. Clean fastq files were generated by filtering out adaptors, contaminants, and low-quality reads (Phred+33 encoding).

After quality control with FastQC, 150bp paired-end reads were aligned to the human reference genome (hg38) using STAR (ref. [8]) with parameters: -alignIntronMin 20 -alignIntronMax 1000000 -alignSJoverhangMin 8 -quantMode GeneCounts”. Gene-level quantification was performed with RSEM (ref. [9]). Data normalization was conducted using the TMM method in edgeR [10]. Principal component and scatter plot analyses assessed data quality post-quantification. Differentially expressed genes were identified based on TMM-normalized counts, with multiple testing correction using the Benjamini-Hochberg procedure [11].

### RNA extraction and RT-qPCR

Total RNA was extracted by direct lysis using the SingleShot Cell Lysis Kit (Bio-Rad, Hercules, CA, USA, #1725080), following the manufacturer’s instructions. Cells were washed once with phosphate buffered saline, lysed in 50 µL of lysis buffer, and incubated at room temperature for 5 min. Lysates (1-2 µL) were used for complementary DNA synthesis with the iScript Reverse Transcription SuperMix (Bio-Rad, #1708841). RT-qPCR was performed using the SsoAdvanced Universal SYBR Green Supermix (Bio-Rad, #1725272) and gene-specific primers on a CFX Opus 96 Real-Time PCR system (Bio-Rad, #12011319). Each reaction was run in triplicates with 200 nM primers. Relative gene expression was analyzed using the 2^-ΔΔCt^ method and normalized to GAPDH or ACTB.

### Protein extraction and western blotting

Proteins were extracted from cells using RIPA buffer (Boston Bioproducts, Milford, MA, USA, #BP-115) supplemented with a Halt protease inhibitor cocktail (Thermo Fisher Scientific, Waltham, MA, USA, #87786). After adding Laemmli SDS sample buffer, reducing (Thermo, #J61337), proteins were reduced and denatured at 95°C for 8 min. Proteins were then separated using Mini-PROTEAN TGX Precast Protein Gels (Bio-Rad, #4568096) and transferred onto nitrocellulose membranes with the Trans-Blot Turbo transfer system (Bio-Rad, #1704150) and a Turbo RTA Mini 0.2 µm nitrocellulose transfer kit (Bio-Rad, #1704270). The membranes were blocked with EveryBlot blocking buffer (Bio-Rad, #12010020) for 5 min and incubated with primary antibodies for 1 h at room temperature (RT) or overnight in a cold room: RIPK2 (Cell Signaling Technology [CST], Danvers, MA, #4142), β-actin (CST, #4970S), JAK1 (CST, #3344T), JAK2 (CST, #3230T), pSTAT3 (CST, #9145T), STAT3 (CST, $12640T), and GAPDH (CST, 51332S) at 1:1 000 dilution. Following washes, the membranes were incubated for 1 h at RT with an anti-Rabbit IgG HRP-linked secondary antibody (CST, #7074S) at a 1:5 000 dilution. After a 5-min incubation with SuperSignal West Pico PLUS chemiluminescent substate (Thermo Fisher, #34580) at RT, signals were visualized using a ChemiDoc MP imaging system (Bio-Rad, #12003154).

### Calculation of RIPK2 signature scores

Signature scores were computed using R (v4.3.1). For qPCR-based analyses, ΔCt values—which are inversely proportional to gene expression levels—were Z-score normalized across samples for each of the eight signature genes. To construct a composite signature score for each biological sample, the Z-scores of RIPK2-repressed genes were summed directly, while the Z-scores of RIPK2-dependent genes and RIPK2 itself were inverted (multiplied by -1) before summation. Finally, the summed Z scores were divided by 8 (the number of signature genes) to facilitate interpretation.

For clinical transcriptomic data, datasets were retrieved from cBioPortal and the Gene Expression Omnibus [12, 13]. Only datasets containing RIPK2 and the eight signature genes were included for further analysis. After removing rows with NA values, log2-transformed expression values were center-normalized. In the resulting expression matrix, RIPK2-repressed signature genes (*i.e.*, LAMB3, PEAR1, PLXNA2, and TRPV4) were assigned a weight of -1. Finally, single-sample Gene Set Enrichment Analysis (ssGSEA) (ref. [14]) was performed to compute signature scores using a nine-gene set comprising RIPK2 and the eight signature genes.

### Survival analysis

Kaplan-Meier survival analysis was performed using the *survival* and *survminer* libraries in R (v4.3.1). The surv_cutpoint() function was used to compute optimal cut points, the survfit() function to generate survival curves, the coxph() function to calculate hazard ratios, and the surv_pvalue() function to compute log-rank *p* values. The ggsurvplot() was used to visualize survival curves. Additionally, the concordance index (*c*-index) was calculated using the concordance.index() function from the *survcomp* library.

### Statistical analysis

Statistical analyses were performed using EXCEL and R (v4.3.1). For cell-based experiments, three biological replicates were conducted unless otherwise specified. Differences between two groups were assessed using unpaired two-tailed Student’s *t*-test. *P* values < 0.05 were considered statistically significance (\**p* < 0.05, \*\**p* < 0.01, \*\*\**p* < 0.001, and \*\*\*\**p* < 0.0001).

## RESULTS

### Identification of candidate RIPK2 signature genes in PC cells

We previously generated CRISPR/Cas9-mediated RIPK2-knockout (*RIPK2*-KO) clones of 22Rv1, DU145, and PC3 cells, which were derived from distinct PC patients, using two independent guide RNAs (gRNAs) [2]. *RIPK2*-KO was confirmed by western blotting (Fig. S1), after which we performed RNA-Seq analysis to identify RIPK2-regulated genes.

Applying a threshold of adjusted *p*-value <0.01 and absolute fold change >1.5, we identified thousands of significantly upregulated or downregulated genes in each *RIPK2*-KO clone compared to their respective control cells (Fig. 1A, Table S1). The exception was the DU145 *RIPK2*-gRNA2 (Rg2) clone, which exhibited 464 significantly upregulated and 945 significantly downregulated genes (Fig. 1A).

**Fig 1.**
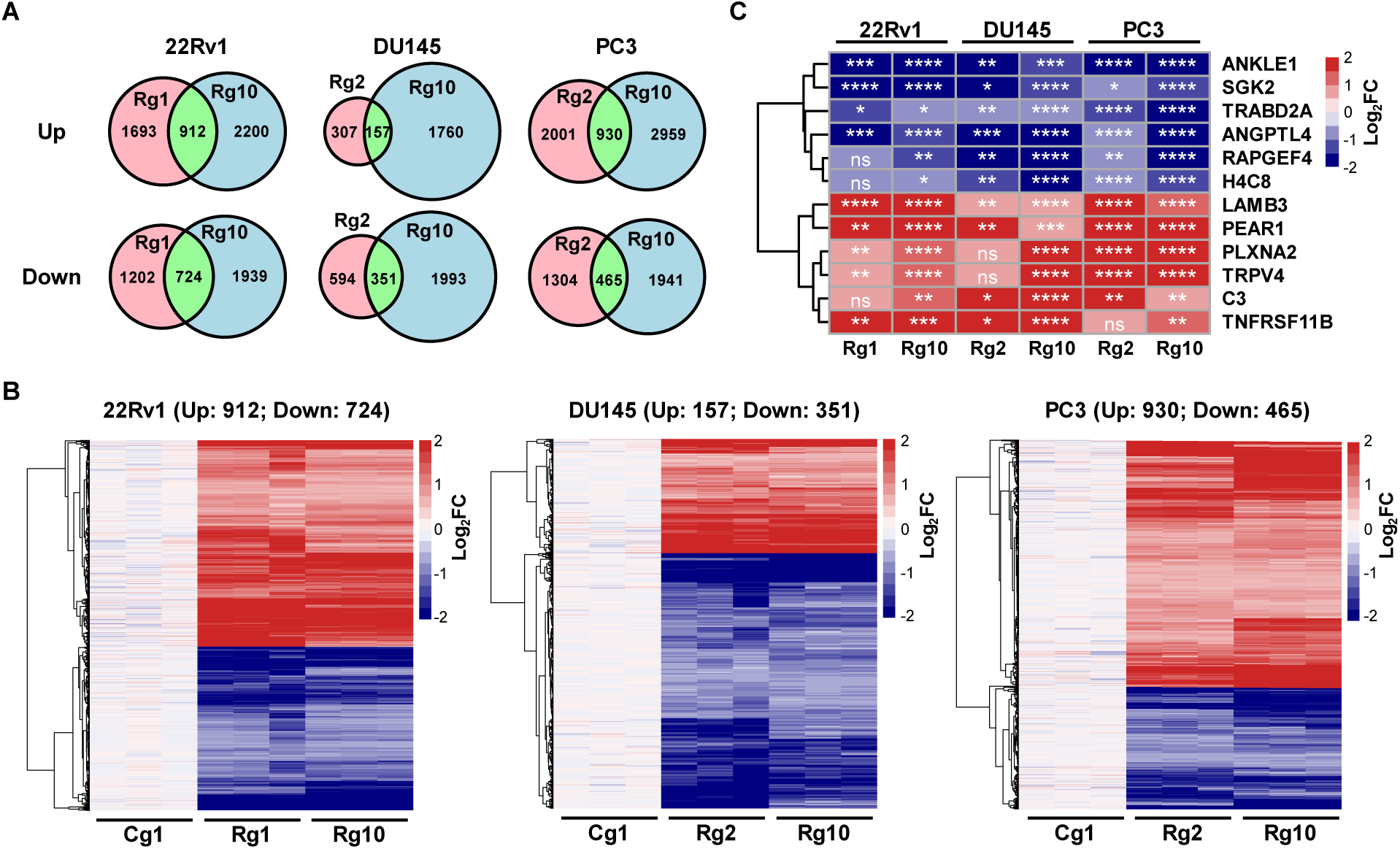
Identification of candidate RIPK2 signature genes in PC cells. **A** Venn diagrams of genes significantly upregulated (top) and downregulated (bottom) by *RIPK2* knockout (KO) in three PC cell lines. Cg1: control gRNA #1; Rg1: RIPK2 gRNA #1; Rg2: RIPK2 gRNA #2; Rg10: RIPK2 gRNA #10. **B** Heatmaps of overlapping genes in different PC cell lines. **C** Heatmap of 13 candidate RIPK2 signature genes. Data were normalized against their respective control cells. Nominal *p*-values were determined by unpaired two-tailed Student’s *t*-test, comparing *RIPK2*-KO to the Cg1 condition. *: *p* < 0.05; **: *p* < 0.01; ***: *p* < 0.001; ****: *p* < 0.0001; ns: not significant; Log2FC: log2-transformed fold change.

Overlap analysis identified a subset of genes consistently upregulated or downregulated in both *RIPK2*-KO clones within each cell line. The differential expression patterns of these genes are illustrated in heatmaps (Fig. 1B).

Recognizing the importance of expression magnitude for biomarker identification, we focused on genes consistently exhibiting an absolute fold change >1.5 across all *RIPK2*-KO clones. This analysis identified 13 candidate RIPK2 signature genes significantly differentially expressed in *RIPK2*-KO cells, with at most one exception among the six *RIPK2*-KO clones (Fig. 1C).

### RT-qPCR validation of candidate RIPK2 signature genes

To validate the candidate RIPK2 signature genes, we performed RT-qPCR analysis on PC3 and 22Rv1 cells using the primers listed in Table S2. For 22Rv1 cells, the Rg2 clone was replaced with the Rg3 clone due to its higher knockout efficiency (Fig. 2A).

**Fig 2.**
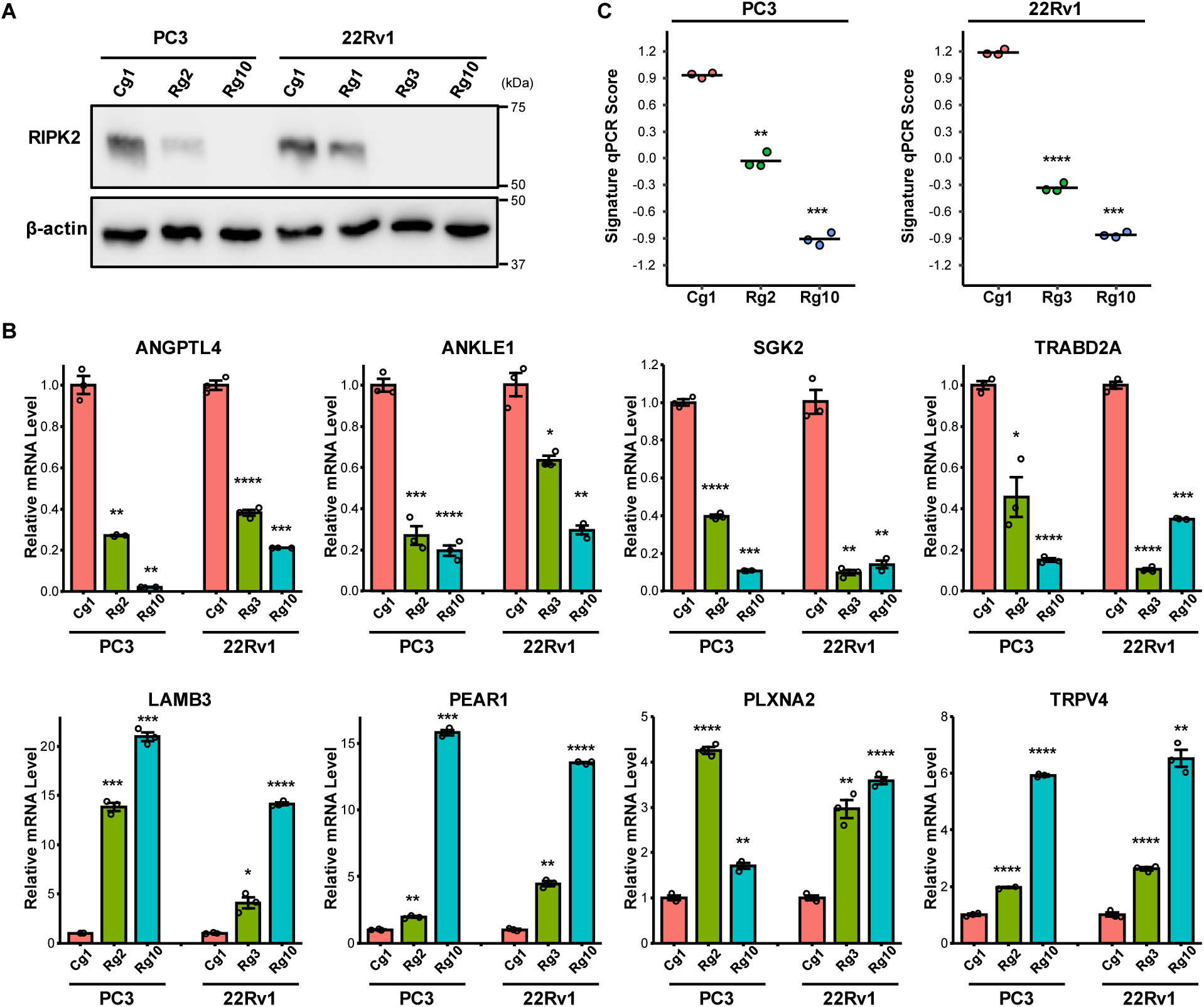
RT-qPCR validation of candidate RIPK2 signature genes. **A** Representative immunoblotting images of the indicated proteins in control and *RIPK2*-KO cells. **B** Bar graphs showing the relative mRNA expression levels of RIPK2-dependent genes (top) and RIPK2-repressed genes (bottom). **C** Swarm plots of RIPK2 signature qPCR scores in control and *RIPK2*-KO cells. The solid lines indicate average qPCR scores. Nominal *p*-values were determined by unpaired two-tailed Student’s t-test. *: *p* < 0.05; **: *p* < 0.01; ***: *p* < 0.001; ****: *p* < 0.0001.

RT-qPCR confirmed the differential expression of eight candidate genes, including four RIPK2-dependent genes, whose mRNA levels were reduced by *RIPK2*-KO, and four RIPK2-repressed genes, whose mRNA levels were increased by *RIPK2*-KO (Fig. 2B). Five additional candidate genes did not show consistent expression patterns and were thus excluded from further analysis (Fig. S2).

Using the eight validated genes, we calculated RIPK2 signature scores for each clone. Consistent with RIPK2 protein levels (Fig. 2A), *RIPK2*-KO clones (Rg2/3 and Rg10) exhibited significantly reduced RIPK2 signature scores, with Rg10 showing the most pronounced reduction (Fig. 2C).

### RIPK2 inhibition modulates signature gene expression and reduces signature scores in a dose- and/or time-dependent manner

To determine whether the RIPK2-regulated gene signature can serve as a marker of RIPK2 activity in PC cells, we treated PC3 and 22Rv1 cells with two distinct RIPK2 inhibitors: ponatinib and RIPK2-IN-4 (RIN4). Ponatinib, a multi-kinase inhibitor clinically approved for leukemia treatment, effectively inactivates the RIPK2/MKK7/c-Myc signaling pathway and reduces PC metastasis *in vivo* [2]. RIPK2-IN-4 is a highly selective and potent RIPK2 inhibitor [15].

Dose-response experiments showed that both inhibitors significantly modulated the expression of signature genes and reduced signature scores in a dose-dependent manner (Fig. 3A-B, Fig. S3). Time-course experiments revealed distinct dynamic responses among individual signature genes following RIPK2 inhibition (Fig. 3C, Fig. S4). Despite these variations, both inhibitors led to a significant, time-dependent decrease in RIPK2 signature scores (Fig. 3D).

**Fig 3.**
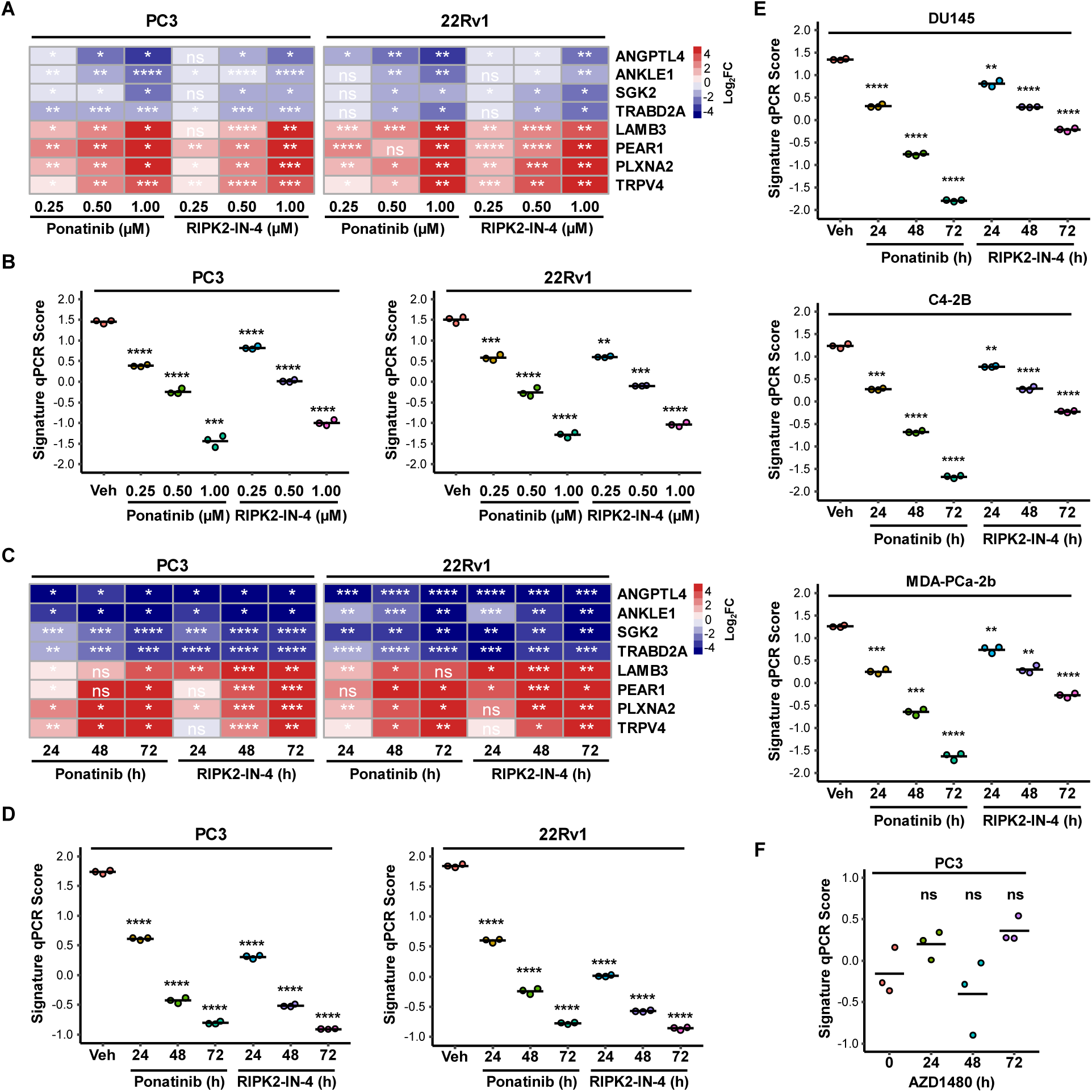
RIPK2 inhibition modulates signature gene expression and reduces signature scores in a dose- and time-dependent manner. **A** Heatmaps of relative mRNA levels of the indicated signature genes in response to the indicated doses of ponatinib and RIPK2 inhibitor 4 (RIPK2-IN-4) for 24 h. Data were normalized against their respective control samples. **B** Swarm plots of signature qPCR scores in response to the indicated doses of ponatinib and RIPK2-IN-4 for 24 h. **C** Heatmaps of relative mRNA levels of the indicated signature genes in response to 1 µM ponatinib and RIPK2-IN-4 for the indicated time points. Data were normalized against their respective control samples. **D** Swarm plots of signature qPCR scores in response to 1 µM ponatinib and RIPK2-IN-4 for the indicated time points. **E** Swarm plots of signature qPCR scores in three additional PC cell lines in response to 1 µM ponatinib and RIPK2-IN-4 for the indicated time points. **F** Swarm plot of signature qPCR scores in PC3 cells in response to 1 µM AZD1480 for the indicated time points. Nominal *p*-values were determined by unpaired two-tailed Student’s *t*-test. *: *p* < 0.05; **: *p* < 0.01; ***: *p* < 0.001; ****: *p* < 0.0001; ns: not significant.

Additionally, both inhibitors significantly modulated the mRNA levels of RIPK2 signature genes and reduced signature scores in DU145, C4-2B, and MDA-PCa-2b cells in a time-dependent manner, with ponatinib exhibiting greater potency than RIPK2-IN-4 in these cell lines (Fig. 3E, Fig. S5). This time dependency was not observed in cells treated with vehicle control (0.1% DMSO) alone (Fig. S6), supporting that the observed changes were due to RIPK2 inhibition. The magnitude of changes in RIPK2 signature genes was more modest in DU145, C4-2B, and MDA-PCa-2b cells than in PC3 and 22Rv1 cells (Fig. S4 versus Fig. S5), possibly because basal RIPK2 activity levels are lower in the former three cell lines than in the latter two.

To exclude the possibility that the RIPK2 signature genes respond to kinase inhibitors in general, PC3 cells were treated with the well-characterized JAK inhibitor AZD1480. Although AZD1480 reduced STAT3 phosphorylation levels in a time-dependent manner, it did not significantly affect RIPK2 signature gene expression or activity scores (Fig. 3F, Fig. S7). Collectively, these findings suggest that the eight signature genes constitute a candidate biomarker for RIPK2 activity in PC cells.

### RIPK2 signature scores correlate more strongly than RIPK2 mRNA levels with poor prognosis in PC

To determine whether RIPK2 signature scores outperform RIPK2 mRNA levels in prognostic value for PC, we incorporated RIPK2 and the eight validated signature genes into a new gene signature and assessed its prognostic significance.

Our analysis revealed that RIPK2 signature scores were significantly higher in metastatic PC compared to primary PC (Fig. 4A). In contrast, RIPK2 mRNA levels did not differ significantly between primary and metastatic PC tissue specimens (Fig. 4A).

**Fig 4.**
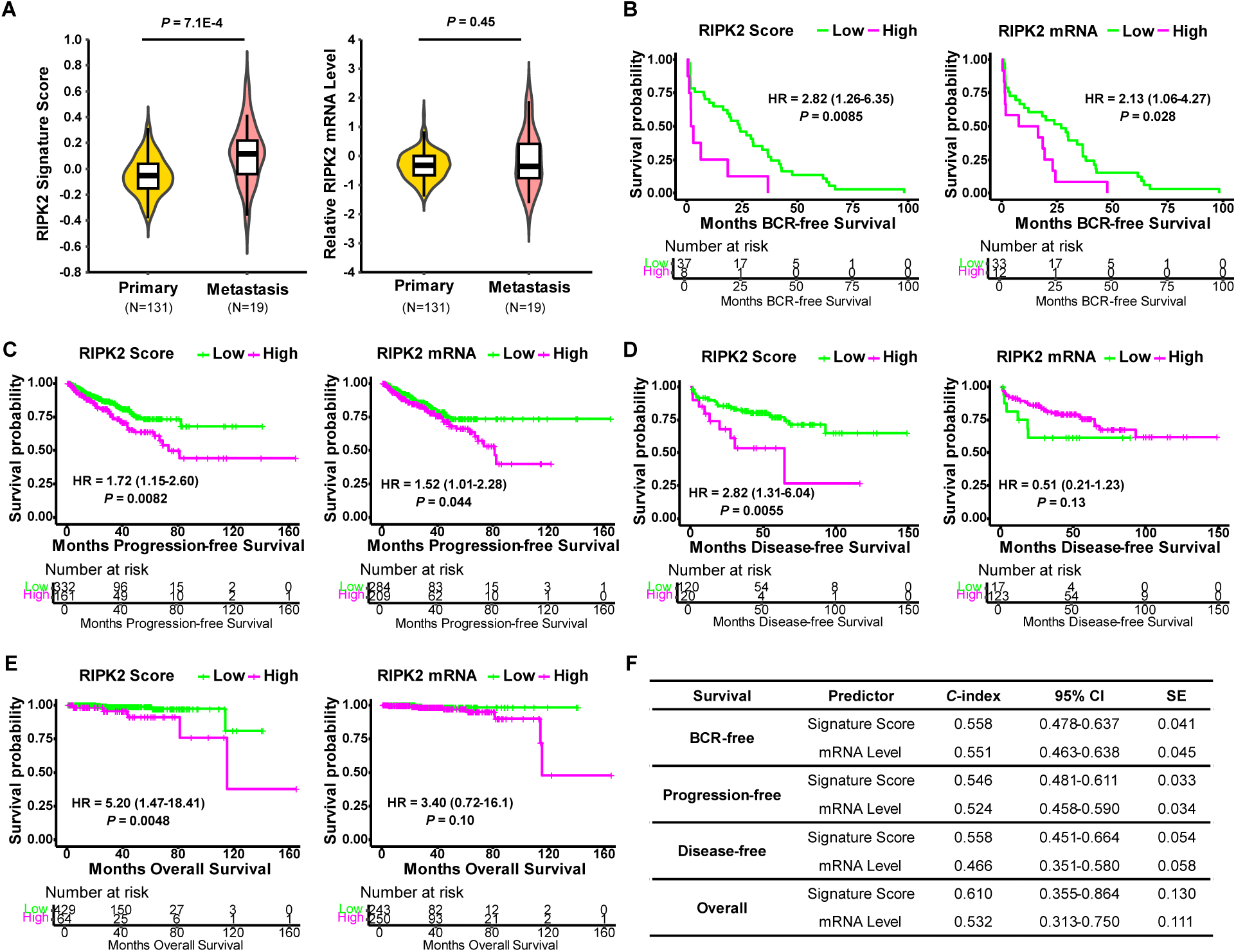
RIPK2 signature scores correlate more strongly than RIPK2 mRNA levels with poor prognosis in PC. **A** Violin plots of RIPK2 signature scores (left) and RIPK2 mRNA levels (right) in primary versus metastatic PC in the Taylor cohort [24]. **B** Kaplan-Meier curves of biochemical recurrence (BCR)-free survival in the GSE70769 cohort [25]. **C** Kaplan-Meier curves of progression-free survival in the TCGA Pan-Cancer Atlas PC cohort. **D** Kaplan-Meier curves of disease-free survival in the Taylor cohort. **E** Kaplan-Meier curves of overall survival in the TCGA Pan-Cancer Atlas PC cohort. **F** Comparison of the prognostic significance of RIPK2 signature scores and mRNA levels. C-index: concordance index; CI: confidential interval; SE: standard error of the estimate. The optimal cut points were used in all Kaplan-Meier curves. HR: hazard ratio (95% confidence interval). Nominal *p*-values were determined using two-sided log-rank test.

Furthermore, RIPK2 signature scores showed stronger correlations than RIPK2 mRNA levels with biochemical recurrence (BCR)-free, progression-free, disease-free, and overall survival in PC, regardless of whether independent optimal cut points (Fig. 4B-E) or the same proportion of patients (Fig. S8) were used.

To enable a cut point-independent comparison between RIPK2 signature scores and RIPK2 mRNA levels, we evaluated their concordance index (*c*-index) values. Consistently, RIPK2 signature scores exhibited higher *c*-index values than RIPK2 mRNA levels for BCR-free, progression-free, disease-free, and overall survival (Fig. 4F).

## DISCUSSION

RIPK2 has recently emerged as a promising drug target in multiple cancers, including PC. However, the absence of reliable biomarkers to measure RIPK2 activity has posed a significant barrier to the development of precision treatment strategies. In this study, we identified and validated an eight-gene signature whose expression levels are consistently modulated by both genetic and pharmacological inhibition of RIPK2. Importantly, RIPK2 signature scores decreased in a dose- and time-dependent manner upon treatment with RIPK2 inhibitors, highlighting their potential utility for monitoring patient responses to anti-RIPK2 therapy. Furthermore, RIPK2 signature scores outperform RIPK2 mRNA levels in prognostic significance, reinforcing their potential clinical relevance.

Although the clinical associations are promising, the biological functions and transcriptional regulation of the RIPK2 signature genes remain largely unexplored. Among the four RIPK2-dependent signature genes, *ANGPTL4* encodes angiopoietin-related protein 4, a multifunctional protein involved in glucose and lipid metabolism, inflammation, angiogenesis, and tumorigenesis [16]. *ANGPTL4* is upregulated in response to hypoxia and promotes PC progression by activating the PI3K/Akt pathway, with high expression levels correlating with shorter BCR-free survival [17]. It also interacts with IQGAP1 at the PC cell membrane, activating the Raf/MEK/ERK/PGC1α pathway to enhance mitochondrial biogenesis and oxidative phosphorylation, ultimately driving PC growth and chemoresistance [18]. *SGK2* encodes serum/glucocorticoid regulated kinase 2, which has been implicated in PC metastasis by inhibiting ferroptosis through upregulation of GPX4 [19]. In comparison, little is known about the roles of *ANKLE1* and *TRABD2A* in PC.

Among the four RIPK2-repressed genes, *PLXNA2* encodes plexin-A2, a coreceptor for SEMA3A and SEMA6A. One study reported that *PLXNA2* contributes to TMRPSS2:ERG-enhanced PC3 cell migration and invasion [20]. *LAMB3*, encoding laminin subunit β3, is a component of laminin 5, a key adhesive protein in the extracellular matrix that regulates cell proliferation, migration, and cell cycle progression in various diseases [21]. *PEAR1* encodes platelet endothelial aggregation receptor 1, a tyrosine kinase-like transmembrane receptor essential for platelet and megakaryocyte biology [22]. *TRPV4* encodes transient receptor potential cation channel subfamily V member 4, a non-selective calcium permeant cation channel involved in osmotic sensitivity and mechanosensitivity [23]. However, the specific contributions of *LAMB3*, *PEAR1*, and *TRPV4* to PC biology remain to be elucidated.

Our findings indicate that the RIPK2 signature is strongly associated with overall survival in PC, with a hazard ratio of 5.2, while its association with BCR-free, progression-free, and disease-free survival is more modest (hazard ratio of 1.7-2.8). This pattern suggests that the RIPK2 signature is not merely a predictor of early disease progression but rather a marker of tumor aggressiveness, metastatic potential, and/or therapy resistance. Given that BCR does not always lead to metastasis or mortality, the stronger correlation with overall survival underscores its potential utility in refining risk stratification beyond traditional prognostic markers. Specifically, this signature may help distinguish patients who require early systemic intervention from those who can be managed with standard-of-care approaches. Further validation in independent cohorts, as well as mechanistic studies, will be critical to fully establish the signature’s clinical utility and to support its integration into precision oncology strategies.

One limitation of our current approach is the reliance on RT-qPCR-based signature scores calculated using Z scores, which are inherently relative, batch-dependent, and sensitive to outliers. Z score normalization assumes a normal distribution of gene expression, compresses biological variability, and reduces direct biological interpretability. To enhance assay consistency and clinical applicability, future efforts will focus on adopting droplet digital PCR to enable absolute quantification of each signature gene, coupled with the development and validation of a fixed scoring algorithm that would allow for standardized comparisons across different tissue specimens and batches.

In conclusion, this study identifies the RIPK2-regulated gene signature as a novel functional biomarker of RIPK2 activity and prognosis in PC, providing a new framework for patient stratification and treatment monitoring. These findings support further clinical validation and suggest potential applications in companion diagnostics for RIPK2-targeted therapies, not only in advanced PC but possibly in other RIPK2-driven cancers. Future studies are warranted to validate the RIPK2 signature in patient-derived tissues and to explore its feasibility in minimally invasive diagnostic platforms, such as liquid biopsy-based assays, for real-time disease monitoring and therapeutic decision-making.

## ACKNOWLEDGEMENTS

We thank Tullia Champney and Lili Guerra for their assistance in lab management. This work was supported by the Department of Defense (DoD) Prostate Cancer Research Program Translational Science Award (W81XWH-22-1-1031), the National Cancer Institute (NCI) R01CA26694, and a startup fund from Stony Brook University (to WY).

## AUTHOR CONTRIBUTIONS

WY conceived and directed the project and calculated the signature scores. AE performed cell culture, drug treatments, RT-qPCR, western blotting, and data analysis. QY and SY analyzed the RNA-Seq data and conducted differential gene expression analysis. ZH performed cell culture, RNA extraction for RNA-Seq analysis, and western blotting. LR conducted cell culture and western blotting. AE and WY wrote the manuscript, with all authors contributing to the writing and providing feedback.

## DATA AVAILABILITY

The datasets generated and/or analyzed in this study are available from the corresponding author upon request. RNA-Seq data have been deposited in the Gene Expression Omnibus (GEO) and are publicly available (GSE292278).

