## Supplementary Figures for "Identification of a RIPK2-Regulated Gene Signature as a Candidate Biomarker for RIPK2 Activity and Prognosis in Prostate Cancer"

### SUPPLMENETARY FIGURES

#### Supplementary Figure S1

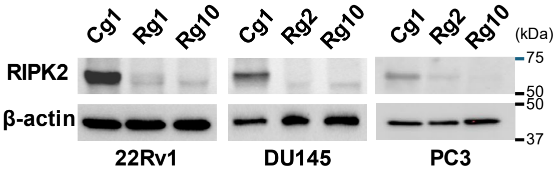


**Supplementary Figure S1. Representative immunoblots of RIPK2 knockout in PC 22Rv1, DU145, and PC3 cells.** Cg1: control gRNA #1; Rg1: RIPK2 gRNA #1; Rg2: RIPK2 gRNA #2; Rg10: RIPK2 gRNA #10.

#### Supplementary Figure S2

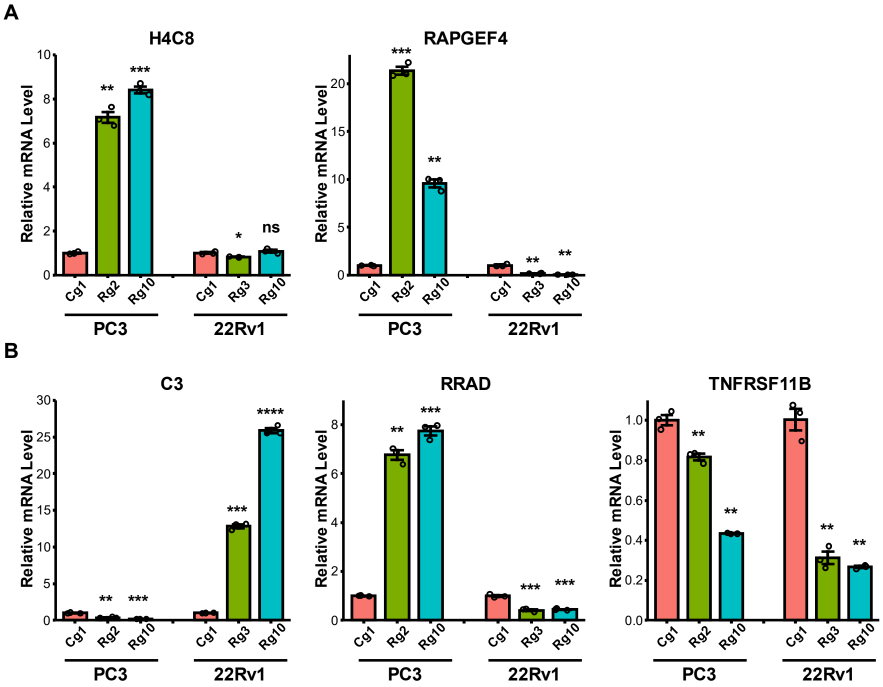


**Supplementary Figure S2. Bar graphs of RT-qPCR results for invalidated candidate signature genes. A** Quantification of the relative mRNA expression levels of the invalidated RIPK2-dependent genes (*i.e.*, genes whose expression decreases upon RIPK2 knockout). **B** Quantification of the relative mRNA expression levels of the invalidated RIPK2-repressed genes (*i.e.*, genes whose expression increases upon RIPK2 knockout). Cg1: control guide RNA #1; Rg2: RIPK2 guide RNA #2; Rg3: RIPK2 guide RNA #3; Rg10: RIPK2 guide RNA #10. Nominal *p*-values were determined by unpaired two-tailed Student's *t*-test, comparing *RIPK2*-KO to the Cg1 condition. Data are presented as mean ± SEM (standard error of the mean). *: $p < 0.05$; **: $p < 0.01$; ***: $p < 0.001$; ****: $p < 0.0001$; ns: not significant.

### Supplementary Figure S3

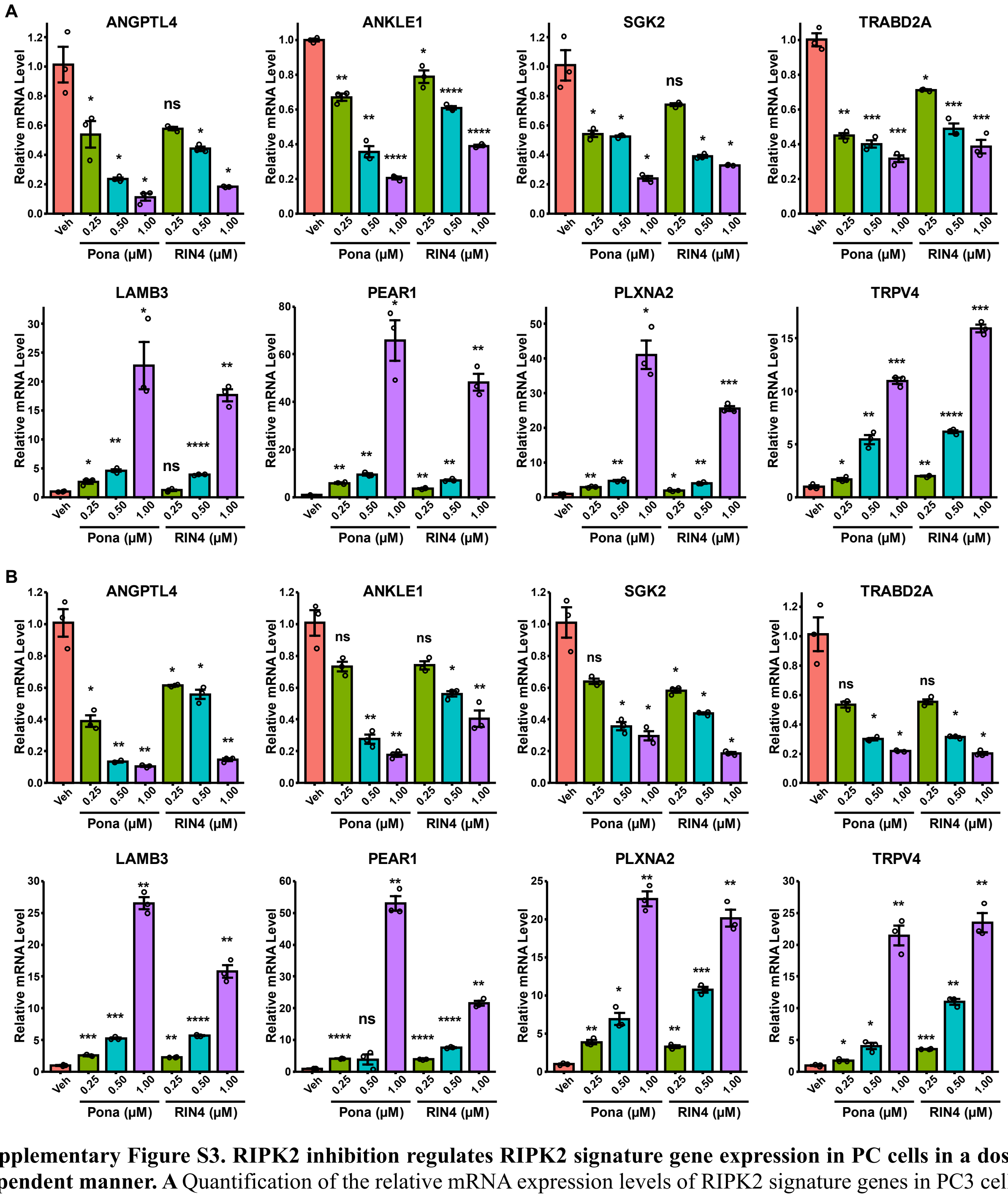


**Supplementary Figure S3. RIPK2 inhibition regulates RIPK2 signature gene expression in PC cells in a dose-dependent manner. A** Quantification of the relative mRNA expression levels of RIPK2 signature genes in PC3 cells. **B** Quantification of the relative mRNA expression levels of RIPK2 signature genes in 22Rv1 cells. Cells were treated with the indicated concentrations of ponatinib (Pona) or RIPK2 inhibitor 4 (RIN4) or with vehicle (Veh) control for 24 h. Nominal $p$-values were determined by unpaired two-tailed Student's $t$-test, comparing each time point to the vehicle control condition. Data are presented as mean ± SEM. *: $p < 0.05$; **: $p < 0.01$; ***: $p < 0.001$; ****: $p < 0.0001$; ns: not significant.

### Supplementary Figure S4

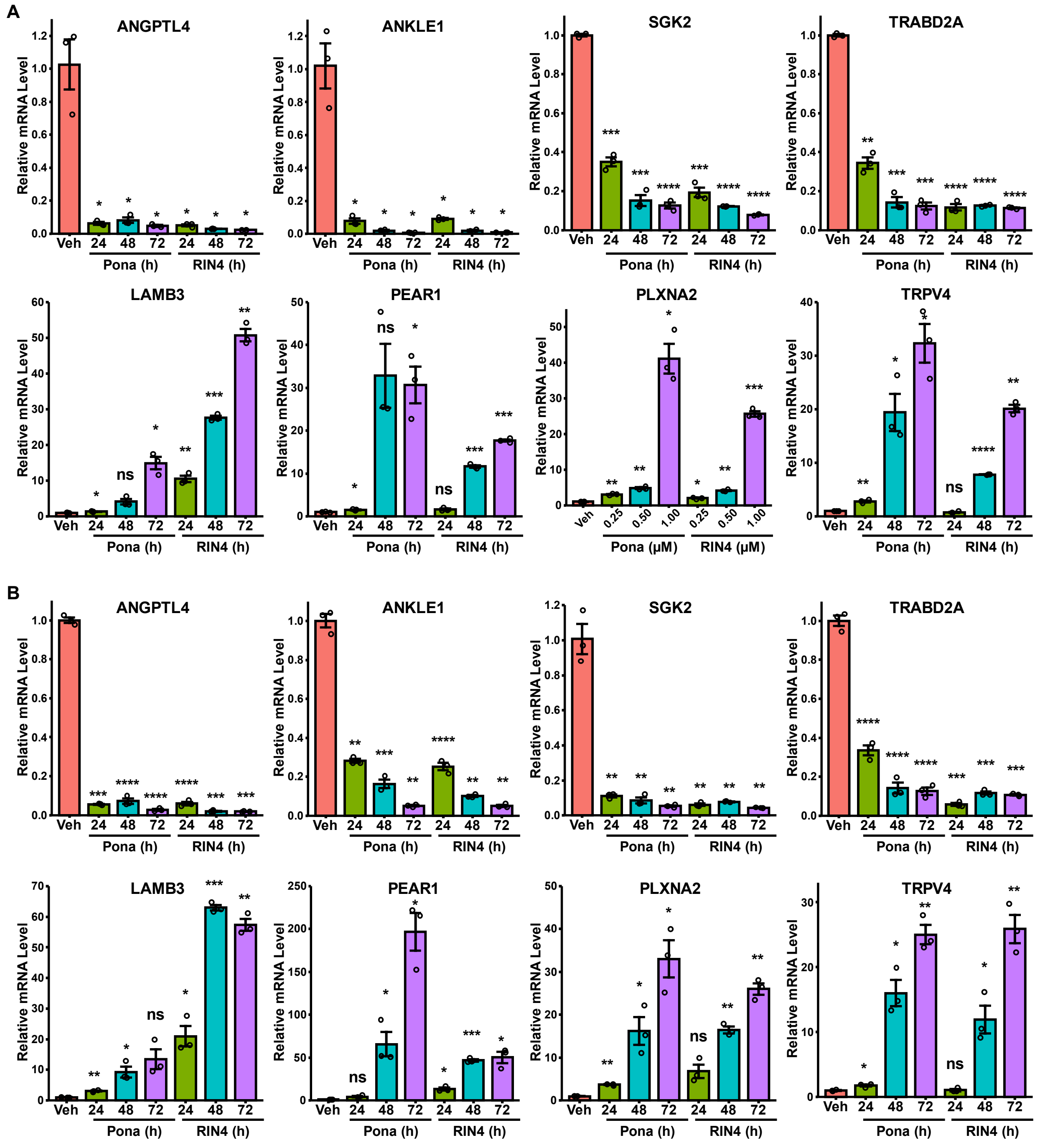


**Supplementary Figure S4. RIPK2 signature genes exhibit different dynamics upon RIPK2 inhibition. A** Quantification of the relative mRNA expression levels of RIPK2 signature genes in PC3 cells. **B** Quantification of the relative mRNA expression levels of RIPK2 signature genes in 22Rv1 cells. Cells were treated with 1 μM ponatinib (Pona) or RIPK2 inhibitor 4 (RIN4) for the indicated time points, or with vehicle (Veh) control for 24 h. Nominal *p*-values were determined by unpaired two-tailed Student's *t*-test, comparing each time point to the vehicle control condition. Data are presented as mean ± SEM. *: $p < 0.05$; **: $p < 0.01$; ***: $p < 0.001$; ***: $p < 0.0001$; ns: not significant.

### Supplementary Figure S5

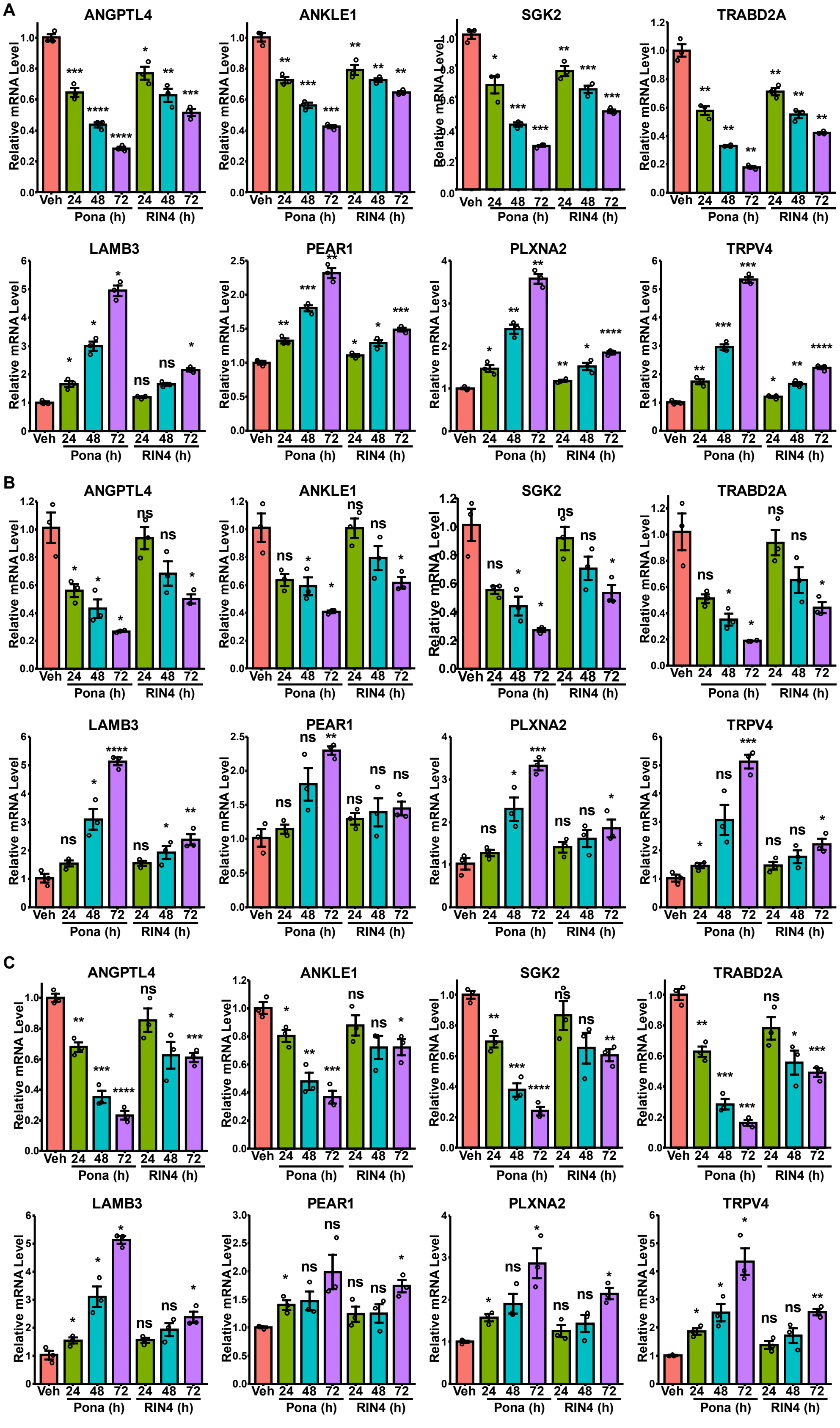

**Supplementary Figure S5. RIPK2 inhibition regulates RIPK2 signature gene expression in three additional PC cell lines in a time-dependent manner. A** Quantification of the relative mRNA expression levels of RIPK2 signature genes in DU145 cells. **B** Quantification of the relative mRNA expression levels of RIPK2 signature genes in C4-2B cells. **C** Quantification of the relative mRNA expression levels of RIPK2 signature genes in MDA-PCa-2b cells. Cells were treated with 1 μM ponatinib (Pona) or RIPK2 inhibitor 4 (RIN4) for the indicated time points, or with vehicle (Veh) control for 24 h. Nominal $p$-values were determined by unpaired two-tailed Student's $t$-test, comparing each time point to the vehicle control condition. Data are presented as mean ± SEM. *: $p < 0.05$; **: $p < 0.01$; ***: $p < 0.001$; ns: not significant.

#### Supplementary Figure S6

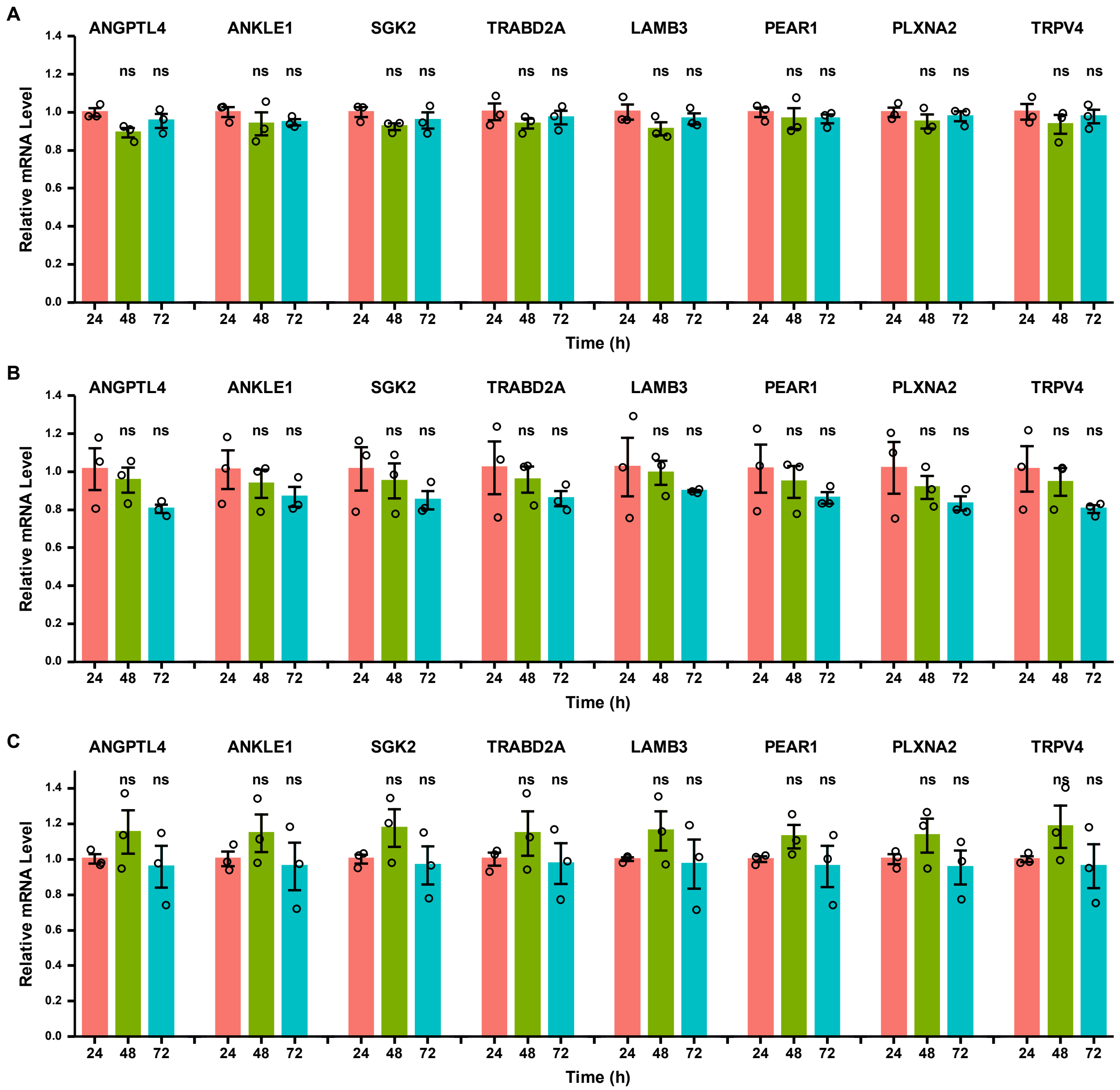


**Supplementary Figure S6. RIPK2 signature gene expression in PC cells is not significantly altered over time under control conditions. A** Quantification of the relative mRNA expression levels of RIPK2 signature genes in DU145 cells. **B** Quantification of the relative mRNA expression levels of RIPK2 signature genes in C4-2B cells. **C** Quantification of the relative mRNA expression levels of RIPK2 signature genes in MDA-PCa-2b cells. Nominal *p*-values were determined by unpaired two-tailed Student's *t*-test, comparing each time point to the 24-h treatment condition. Data are presented as mean ± SEM. ns indicates not significant.

#### Supplementary Figure S7

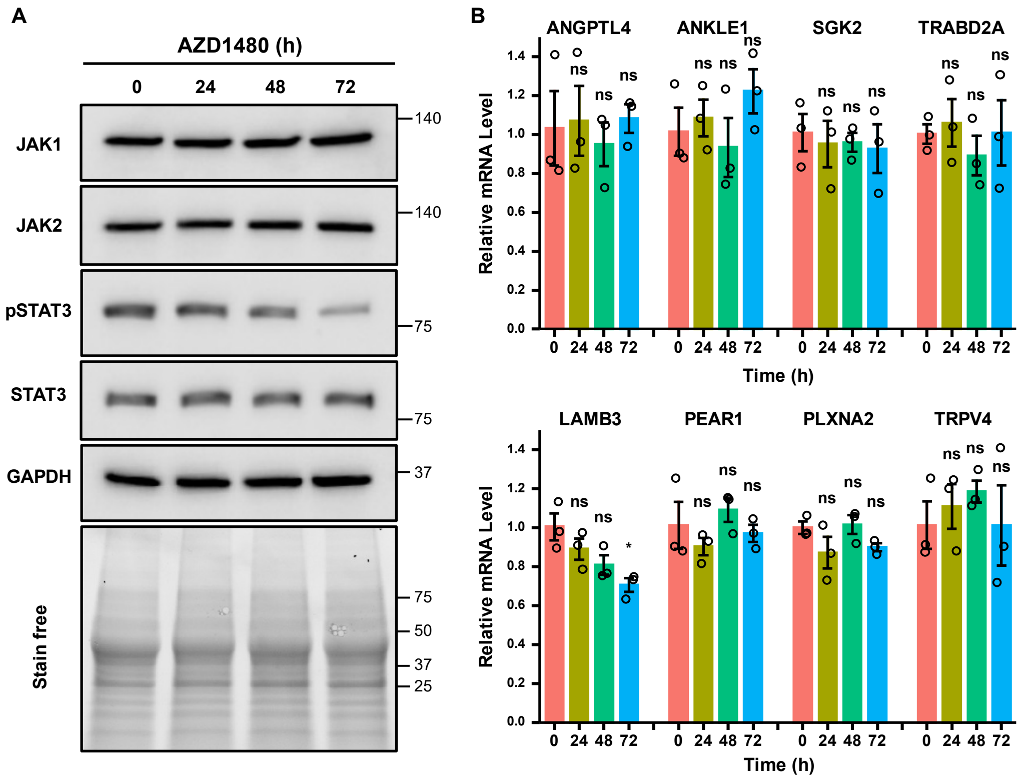


**Supplementary Figure S7. The Janus-associated kinase (JAK) inhibitor AZD1480 has minimal effect on the mRNA expression of RIPK2 signature genes. A** Representative immunoblots of the indicated proteins in total lysates from PC3 cells treated with 1 µM AZD1480 for the indicated time points. **B** Quantification of the relative mRNA expression levels of the indicated RIPK2-dependent (top) and RIPK2-repressed (bottom) genes in PC3 cells treated with 1 µM AZD1480 for the indicated time points. Nominal *p*-values were determined by unpaired two-tailed Student's *t*-test, comparing each time point to the untreated condition. Data are presented as mean ± SEM. * indicates $p < 0.05$; ns indicates not significant.

### Supplementary Figure S8

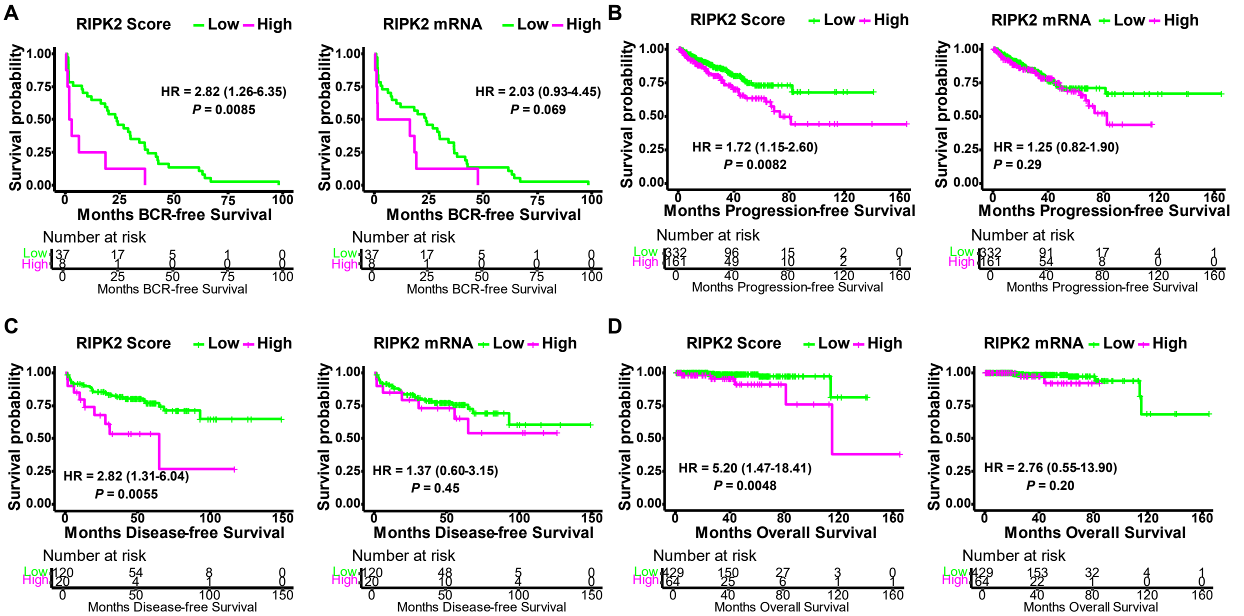


**Supplementary Figure S8. RIPK2 signature scores correlate more strongly with poor prognosis in PC than RIPK2 mRNA levels when stratifying the same proportion of patients. A** Kaplan-Meier curves of BCR-free survival in the GSE70769 cohort. **B** Kaplan-Meier curves of progression-free survival in the TCGA PanCancer Atlas PC cohort. **C** Kaplan-Meier curves of disease-free survival in the Taylor cohort. **D** Kaplan-Meier curves of overall survival in the TCGA PanCancer Atlas PC cohort. Optimal cut points were used for RIPK2 signature score-stratified Kaplan-Meier curves, while the same proportion of patients was used to define cut points for RIPK2 mRNA-stratified Kaplan-Meier curves. HR: hazard ratio (95% confidence interval). Nominal *p*-values were determined using two-sided log-rank test.
